# What can computational modeling tell us about the diversity of odor-capture structures in the Pancrustacea?

**DOI:** 10.1101/337808

**Authors:** Lindsay D. Waldrop, Yanyan He, Shilpa Khatri

**Affiliations:** Author of Correspondence. Dept. of Biology, New Mexico Institute of Mining and Technology, Socorro, NM, USA 87801; Dept. of Mathematics New Mexico Institute of Mining and Technology, Socorro, NM, USA 87801; Applied Mathematics Unit, School of Natural Sciences University of California, Merced Merced, CA 95343.

**Keywords:** olfaction, sensilla, insect, computational modeling, fluid dynamics, sniffing

## Abstract

A major transition in the history of the Pancrustacea was the invasion of several lineages of these animals onto land. We investigated the functional performance of odor-capture organs, antennae with olfactory sensilla arrays, through the use of a computational model of advection and diffusion of odorants to olfactory sensilla while varying three parameters thought to be important to odor capture (Reynolds number, gap-width-to-sensillum-dameter ratio, and angle of the sensilla array with respect to oncoming flow). We also performed a sensitivity analysis on these parameters using uncertainty quantification to analyze their relative contributions to odor-capture performance. The results of this analysis indicate that odor capture in water and in air are fundamentally different. Odor capture in water and leakiness of the array are highly sensitive to Reynolds number and moderately sensitive to angle, whereas odor capture in air is highly sensitive to gap widths between sensilla and moderately sensitive to angle. Leakiness is not a good predictor of odor capture in air, likely due to the relative importance of diffusion to odor transport in air compared to water. We also used the sensitivity analysis to make predictions about morphological and kinematic diversity in extant groups of aquatic and terrestrial crustaceans. Aquatic crustaceans will likely exhibit denser arrays and induce flow within the arrays, whereas terrestrial crustaceans will rely on more sparse arrays with wider gaps and little-to-no animal-induced currents.

## 1 Introduction

### 1.1 Odor capture in the Pancrustacea

Collecting information contained in chemical stimuli, or odors, is a primary way for an animal to interface with its external environment. Animals, including crustaceans and insects, routinely use odors to find food (Rittschof and Sutherland, 1986; Kamio and Derby, 2017; Solari et al, 2017), symbiont hosts (Ambrosio and Brooks, 2011), to recognize individual conspecifics (Gherardi et al, 2005; Gherardi and Tricarico, 2007), to mediate reproduction (Gleeson, 1980), and to avoid predators (Diaz et al, 1999; Pardieck et al, 1999). Odors can illicit behaviors by acting as signals or cues, and these behaviors can be either innate or learned (Derby and Weissburg, 2014).

An important development in the Pancrustacea or Tetraconata, a group including crustaceans and insects, was the invasion of land. Within the Pancrustacea, several lineages evolved independently to live in terrestrial habitats, including Isopoda, Amphipoda, Coenobitidae (terrestrial hermit crabs and *Birgus latro)* (Greenaway, 2003; Hansson et al, 2011; Harzsch and Krieger, 2018). With this change in habitat came a change in the physical properties of the fluid surrounding these animals. Since odor capture and the nature of the odor signal created by environmental flows are both dependent on the physical characteristics of the fluid, the nature of odor capture fundamentally changed between water and air. Has this transition to a terrestrial environment influenced the morphology of odor-capture structures? And can we detect a functionally important signal in the diversity of odor-capture structures that reflect the differences imposed by changing the physical properties in which these structures operate?

### 1.2 Fluid dynamics of odor capture

From odor source to animal, the fluid dynamics of environmental flows create complex odor plumes, discontinuous series of high-concentration odor pulses, that animals must interpret to navigate (Murlis et al, 1992; Weissburg, 2000; Koehl et al, 2001; Dickman et al, 2009; Webster and Weissburg, 2009; Reidenbach and Koehl, 2011). Animals must capture the information contained within these complicated plumes along with fluid-dynamic cues in order to navigate to the source (Moore et al, 1991; Atema, 1995; Page et al, 2011a,b; Weissburg, 2011; Weissburg et al, 2012). Odor capture is the process by which odorants are extracted from the environmental fluid, and it is an important step in olfaction as a whole (Schneider et al, 1998; Kepecs et al, 2006; Moore and Kraus-Epley, 2013). Typically, specialized structures interact with moving fluid, produced by drawing air into a cavity (in the case of mammals) or moving an external chemosensory surface through a fluid (in the case of many crustaceans which use external arrays of hair-like sensilla mounted on antennae).

Odor capture by sensilla arrays depends on the physical interactions between the sensilla array and fluid movements, created by the animals and by environmental flows. Environmental fluid movement, such as wind or water currents, creates odor plumes by dispersing dissolved odorants from the source into the environment. Odorant molecules that dissolve from a source into the surrounding fluid are pulled by turbulent mixing to create high concentration filaments of odor. At large time scales, odors appear to have a Gaussian distribution (averaged across space and time), but at small time and spatial scales (comparable to those experienced by small animals such as insects and other crustaceans) are complicated patterns of odor filaments with varying widths, frequencies, and concentrations (Murlis et al, 1992; Weissburg, 2000; Dickman et al, 2009; Bingman and Moore, 2017). Animals use this information to interpret the location of odor sources (Cardé and Willis, 2008).

The characteristics of an odor plume vary with the properties of the environmental fluid (air or water) and an odorant’s ability to diffuse in that medium. A fluid’s density *(ρ)* and dynamic viscosity (*μ*) will affect the size and frequency of turbulent eddies that create odor filaments at the source, producing relatively larger, less frequent eddies in water than in air (Murlis et al, 1992; Weissburg, 2000; Webster and Weissburg, 2009; Weissburg, 2011). The rate of diffusion of an odorant, quantified by the diffusion coefficient (D), is typically several orders of magnitude smaller in water than in air. This creates odor filaments that are highly concentrated and thin in water (where diffusion is slower) and relatively wider and less concentrated in air (where odorants diffuse more rapidly) (Murlis et al, 1992; Weissburg, 2011). These create distinct patterns of odorant signals at the size and time scale of a sensilla array (Reidenbach and Koehl, 2011; Bingman and Moore, 2017).

In addition to environmental fluid flow, many aquatic and some terrestrial crustaceans and insects waive, or flick, their antennal olfactory arrays to generate fluid movement during odor capture. This fluid movement serves many purposes: it introduces a new sample of fluid close to the sensory structure and moves previously sampled fluid away (Schmidt and Ache, 1979); it thins the attached layer of fluid around the solid sensory structure (the fluid boundary layer) so that molecular diffusion acts over a shorter distance (Stacey et al, 2002; Koehl, 2011); and it increases the temporal and spatial sampling of the fluid environment, thereby increasing the probability of detecting rare or discontinuous odor signals (Kepecs et al, 2006; Koehl, 2006; Cardé and Willis, 2008).

The amount fluid penetration, or leakiness, in an array of sensilla depends on the interactions of the boundary layers around the individual sensillum and the distance between the sensilla. The relative thickness of the boundary layer depends on the Reynolds number *(Re)*:

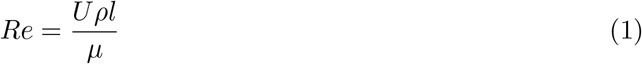

where *l* is a characteristic length scale (such as the diameter of a sensillum) and *U* is the fluid speed relative to the object. Higher flow speeds result in higher *Re* and thinner boundary layers. Cheer and Koehl (1987a,b) described the relationship between *Re* and gap-width-to-sensillum-diameter ratio (*Gw*) between sensilla in the critical ranges that crustacean sensilla arrays occupy (0.01 < *Re* < 10 and *Gw*). When *Re* is low or sensilla are spaced further apart, individual boundary layers do not interact and fluid flow is able to penetrate the array, bringing odor molecules very close to the sensillum surfaces, where diffusion takes odor molecules the final distance to the sensillum’s surface. If flow is slow enough or sensilla are close together, individual boundary layers begin to overlap and drive flow around the array (as opposed to through it), restricting access of odorant molecules to the inner sensory surfaces of the array (Stacey et al, 2002; Schuech et al, 2012).

Air and water differ in terms of both fluid density *(ρ)* and viscosity (*μ*), which will affect the leakiness of the same array in each fluid. Air is 850 times less dense than water and its dynamic viscosity is 59 times lower, leading to *Re* being 15 times lower when an array operates in air as opposed to water. Waldrop and Koehl (2016) calculated that the same antennal array would experience a dramatic decrease in leakiness when moved from water to air that could affect odor-capture performance.

The delivery of odorant molecules to sensory structures depends not just on fluid movement, but also on the rate of diffusion in the fluid. Odor capture also relies on the scaling of advective flows versus the rate of diffusion. The Péclet number (Pe) describes the relative importance of advection (bulk fluid movement) to diffusion:

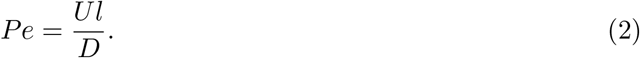

For *Pe* > 1, advection dominates transport of odorant molecules to a structure, whereas for *Pe* < 1, diffusion dominates transport.

Typically, molecules will have much higher diffusion coefficients in air compared to water due to the lower density of air. As a result, *Pe* for the same molecule can be 100 to 10,000 times lower in water compared to air. For a sensillum array similar size to the antennae of marine crustaceans responding to an odorant in air and water, *Pe* can be over 1 in water and below 1 in air (Waldrop and Koehl, 2016; Waldrop et al, 2016). This suggests by simply changing the fluid in which the sensillum array is operating there may be a major shift in the dominant form of mass transport to the array, potentially altering the selective pressures on the array’s morphology (Mellon and Reidenbach, 2012).

### 1.3 Diversity in antennal functional morphology

Many species within the Pancrustacea have sensory structures that consist of external arrays of sensillum-like sensilla concentrated on antennae that protrude away from the head (Fig. 1). The types of sensilla vary, and as a result antennae can provide a range of sensory modalities, including olfaction, gustation, and mechanosensation.

The olfactory hardware of malacostracan crustaceans and insects are similar (Harzsch and Krieger, 2018). The hair-like olfactory sensilla of insects and malacostracan crustaceans consist of a hollow cylindrical tube of cuticle innervated by olfactory sensory neurons, which project outer dendritic segments into the body of each sensillum (Hallberg and Skog, 2011). The cuticle in malacostracan crustaceans is permeable to a variety of chemicals and ions (Gleeson et al, 2000a,b), and the cuticle of insect sensilla is impermeable with pores (Zacharuk, 1980; Keil and Steinbrecht, 1984). Receptors on the outer dendritic segments generate action potentials when exposed to odorants, which are then relayed to the olfactory bulb of the brain. The organization of the olfactory areas of the brain are so similar, Harzsch and Krieger (2018) suggests they reflect a deep homology between malacostracan crustaceans and insects.

The length, diameter, flexibility, number, and arrangement of olfactory sensilla vary widely across the Pancrustacea. Malacostracan crustaceans possess arrays of specialized olfactory sensilla called aesthetascs on their first antennae (antennules) (Fig. 1A-D). Antennules can bear arrays which can range from single lines of short, stiff aesthetascs per segment (Fig. 1 A,C) (Derby, 1982; Grünert and Ache, 1988; Goldman and Patek, 2002) to dense plumes of long aesthetascs (Ghiradella et al, 1968; Snow, 1973; Gleeson, 1980). Coenobitid crabs possess aesthetascs that are shorter, blunter, and more densely packed than their closest relatives (marine hermit crabs) (Ghiradella et al, 1968; Stensmyr et al, 2005; Hansson et al, 2011).

**Figure 1:**
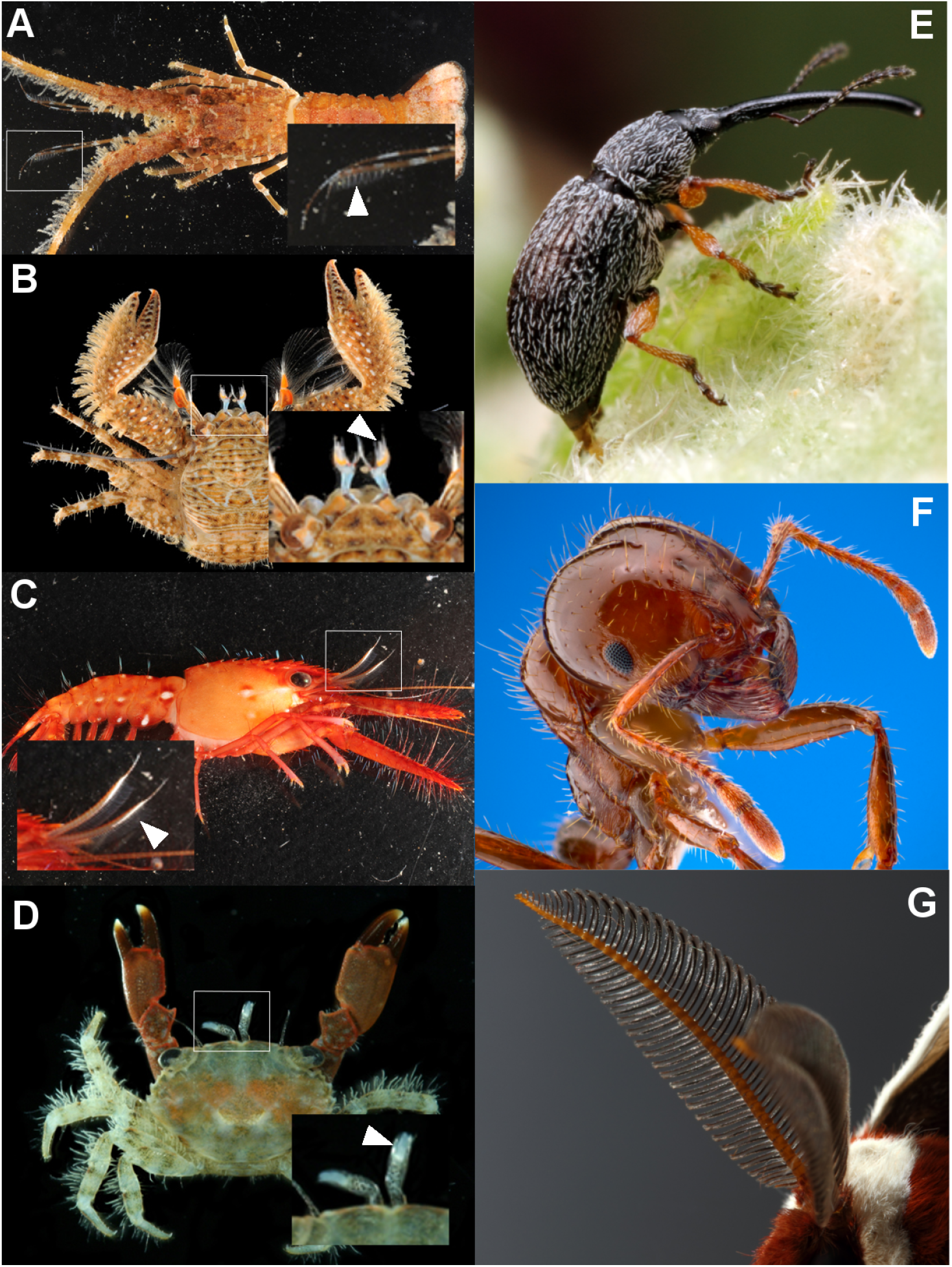
Select antenna morphologies of the Pancrustacea. A–D: aquatic crustaceans (photos courtesy of J. Poupin & the CRUSTA Database (Legall and Poupin, n.d.)), insets highlight first antennae and white arrows indicate aesthetasc arrays. E–F: insects (© Alex Wild, used with permission) with prominent antennae. A: Spiny lobster, *Panulirus argus,* B: banded porcelain crab, *Petrolisthes galathinus,* C: red Hawaiian reef lobster, *Enoplometopus occidentalis,* D: aiyun-tenaga-ohgigani, *Chlorodiella laevissima,* E: hollyhock weevil, *Rhopalapion longirostre,* F: red imported fire ant, *Solenopsis invicta,* G: cecropia silkmoth, *Hyalophora cecropia.*

Insects possess several types of olfactory sensilla, some specialized to specific odorants (Zacharuk, 1980). Antennae can contain one or more types of these sensilla in a variety of different arrangements which can depend on species, sex, and ontogenetic stage (Fig. 1E–F) (Hallberg and Hansson, 1999; Lopez et al, 2014). Arrangements of sensilla range from simple, short arrays of sensilla protruding from the antenna (Fig. 1E,F) to silkmoth antennae which bear dense arrays of specialized sensilla sensitive to sex pheromones (Fig. 1G).

Due to the importance of both fluid movement and diffusion in odor capture, it is unclear to what extent morphological differences in antennal arrays, either within taxa or across taxa, lead to differences in functional performance. Many studies have examined fluid flow through aesthetasc arrays of aquatic and terrestrial malacostracan crustaceans as a proxy for odor capture, focusing on the role of *Re* and *Gw* in sensilla arrays (Mead et al, 1999; Reidenbach et al, 2008; Waldrop et al, 2015a,b). Crayfish exhibit longer aesthetascs with wider gap widths in areas of low flow (Mead, 2008). Terrestrial hermit crabs exhibit a reduction in aesthetasc length and density compared to marine crabs (Ghiradella et al, 1968; Snow, 1973; Mellon and Reidenbach, 2012), and this likely leads to increased odor capture in air (Waldrop et al, 2016). There are other reductions of aesthetasc and brain features in terrestrial or semi-terrestrial brachyruan crabs, making it unlikely that these animals engage in olfaction in air (Krieger et al, 2015). To date, there has been no systematic study of how common features of antennal array morphology affect odor-capture performance across the Pancrustacea.

### 1.4 Computational modeling and evolution

Understanding diversity in sensilla-array morphology and its interaction with the environmental flows in part requires understanding performance over a wide variety of antennal morphologies and flow conditions. There are a large number of parameters associated with the morphology of the arrays and the kinematics of movement, such as the number of sensilla, spacings between sensilla, diameters of sensilla, their position relative to the central support structure. Additionally, there are a large number of parameters based on environmental conditions that will also alter the performance of the array, including *Re* and *Pe* numbers.

Computational models represent a cheap, efficient way at evaluating a relevant performance metric over a very large range of existing and theoretical sensilla-array morphologies. Coupled advection-diffusion studies are better suited to evaluate functional performance of an array during odor capture than examining leakiness alone, since it takes into account the role of diffusion rates into capture. A wide range of morphologies and environmental conditions can be mimicked through modeling and the subsequent impact on odor-capture performance measured.

Variation in these parameter inputs, the raw material on which natural selection works, will affect the performance outputs in complex ways. Altering single variables could have oversized effects on performance or no effects at all. Systems with few degrees of freedom can have several combinations of inputs that produce the same performance (“many-to-one mapping”) (Wainwright, 2007; Anderson and Patek, 2015). Making sense of the holistic effects of input variation on performance output is a key step in determining how functional performance impacts the creation of morphological diversity, despite being often overlooked in many studies (Patek, 2014).

However, the lack of analysis tools that can quantitatively describe parameter effects on computational models limits these models’ ability to inform studies of morphological diversity and evolution. In this study, we use uncertainty quantification to quantify the relative effects of change on performance and provide sensitivity analyses for individual parameter and parameter combinations. Using these sensitivity analyses will allow us to assess which parameters are relatively more sensitive than others and then make specific predictions about what exists in corresponding natural systems. Parameters that are very sensitive to change can either be highly constrained and show little diversity in morphology across clades or be the basis of very fast morphological change within a clade (Anderson and Patek, 2015; Muñoz et al, 2017). Conversely, parameters that are not very sensitive could be free to diversify and show high levels of morphological variation without significant sacrifices to functional performance.

### 1.5 Study Objectives

In this study, we have created a computational model of advection and diffusion to study the impact of variation in morphological parameters on the functional performance of olfactory sensillum arrays in differing fluid environments. This model uses an idealized antennal sensillum array, representing olfactory sensilla, to assess the sensitivity of three morphological and kinematic parameters: the *Re* of the fluid movement relative to the sensilla, the gap-to-diameter ratio of the sensilla (*Gw*), and the angle of the array to the direction of oncoming flow (*θ*). This array is tested in two chemical fluid environments – air and water – using a typical odorant filament and diffusion coefficient characteristic of each environment.

With this model, we will address the following questions:

1. Do features of flow (average speed and shear rates around sensilla, leakiness) predict odor-capture performance?
2. Are there differences in how parameters (*Re*, *Gw, θ*) affect odor-capture performance in air and water?
3. Can sensitivity analyses generate hypotheses to predict patterns of morphological diversity in extant groups, and are these hypotheses different for aquatic and terrestrial groups within the Pancrustacea?

## 2 Materials and Methods

### 2.1 Computational Model

#### 2.1.1 Constraint-based Immersed Body Method

In order to simulate fluid flow around the boundaries of each sensillum and antenna, we used the constraint-based immersed body method (cIB) (Sharma et al, 2005; Bhalla et al, 2013; Kallemov et al, 2016), a version of the regular immersed boundary method (IBM). The IBM, developed by Peskin (Peskin, 2002), fully couples the motion of an elastic boundary with the resulting fluid flow. In the IBM, the incompressible Navier-Stokes equations for fluid flow are solved on a Eulerian grid using an external forcing term (*F*(*x, t*)) modeling the force on the fluid from the Lagrangian boundary:

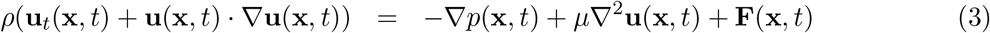

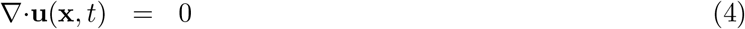

where u(x, *t*) is the fluid velocity, *p*(x, *t*) is the pressure, *ρ* is the fluid density, and *μ* is the dynamic viscosity of the fluid. The independent variables are the time *t* and the position x.

The immersed boundary is modeled using Lagrangian points. The interaction equations between the fluid Eulerian grid and the boundary Lagrangian points are given by:

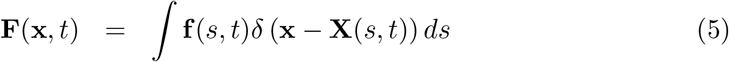

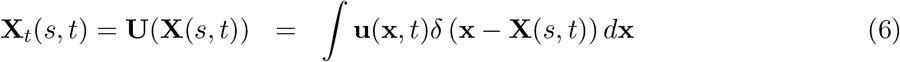

where **X**(*s, t*) gives the Cartesian coordinates at time *t* of the Lagrangian point labeled by parameter s and f(*s,t*) is the force per unit length applied by the boundary to the fluid. In these equations, the two-dimensional delta function, *δ*(x – **X**(*s,t*)), is used to go between the Lagrangian variables and the Eulerian variables. As stated above, eq. 5, gives the force from the boundary on the fluid grid. Eq. 6 gives the velocity of the boundary, **X**_*t*_(*s,t*) = **U**(**X**(*s,t*)), due to the fluid flow.

In the cIB, stead of treating each point separately (as is the case for the regular IBM), the motion of the entire object represented by points is constrained and prescribed. The additional force due to the existence of this body is added to eq. 4 for areas inside the internal volume created by the series of points. The object boundary does not required connections of springs and beams or meshing, making it more computationally efficient.

We used an implementation of this method in the Immersed Boundary with Adaptive Mesh Refinement (IBAMR) package with the constraint IB solver (Bhalla et al, 2013). IBAMR uses local grid refinement to structure the cartesian grid on which the discretized incompressible Navier-Stokes equations are solved, producing a grid that is fine close to the boundary and courser away from the boundary to reduce computational time (see Griffith and Peskin (2005); Griffith (2009); Griffith and Lim (2012) for additional details on IBAMR).

The array was modeled in two dimensions as four solid circles (representing four solid cylinders with a circular cross-sectional area in three dimensions): three smaller cylinders representing olfactory sensilla (diameters, l = 0.01 m) evenly spaced in an array and a fourth representing the supporting antenna (antenna diameter = 0.1 m) (Fig. 2A). This hypothetical array was not modeled after an individual species or group of animals, but represents characteristic features of several groups which were found to have some effects over fluid flow within sensillum arrays, including gap-to-sensillum-diameter ratio (*Gw*) and angle of the array with respect to flow (*θ*) (Cheer and Koehl, 1987b,a; Loudon et al, 2000; Reidenbach et al, 2008; Nelson et al, 2013; Waldrop, 2013; Waldrop et al, 2014, 2015b; Waldrop and Koehl, 2016). The gap between the edge of the center sensillum and the edge of the antenna was maintained at 0.02 m. The spacing between the center of each sensilla that made up the array was determined by a gap-to-diameter ratio that varied between 1.4 and 49. The angle of the array relative to flow (positive x-axis) was varied between 3.57 and 176 degrees. Each cylinder was modeled using evenly spaced points (spaced apart by 4.88 × 10^−4^ m).

**Figure 2:**
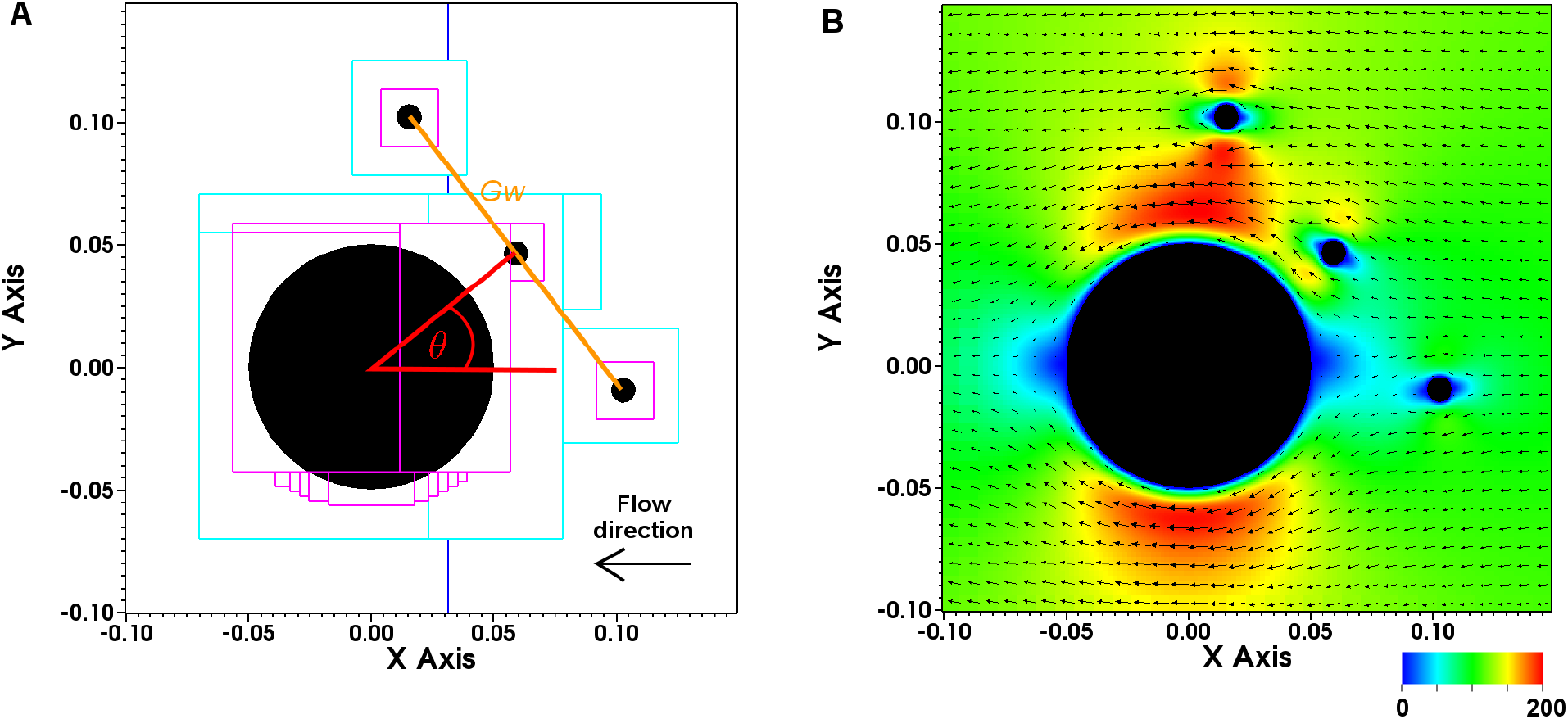
Two-dimensional model of an antennule (large black; circle) and three olfactory sensilla (small blac° circles). A. Flow) directìon is parallel to tire x-axis in the negotive direction (black arrow, right to left). The distance of the array from the antennule was fixed at 0.01 m. Colored boxes (dark blue, light blue, purple) indicate initial levels of the adaptive meshing on the Eulerian fluid grid used to calculate velocities, increasing in fineness. Parameters altered in the study are the angle of the array with respect to flow direction (*θ*, red angle) and the gap-to-sensillum-diameter ratio (*Gw*, orange line). B. A sample velocity field resulting from the simulation of fluid flow through the array (*Re* = 0.24, *Gw* = 6.08, *θ* = 38). Color indicates magnitude of fluid velocity in non-dimensional units and black arrows indicate direction and magnitude of fluid velocity. Plots were generated using VisIt (Childs et al, 2012).

Flow past the sensilla arrays was produced by imposing Dirichlet boundary conditions at the positive and negative x-axis boundaries of the domain as a fixed horizontal flow speed of *U_x_* and 0 for the positive and negative y boundaries. At the beginning of each simulation, *U_x_* was linearly increased from 0 to a final steady-state value to accelerate the flow past the sensilla array. The speed of the flow tank’s ambient flow was quickly accelerated and then set to a constant 0.06 ms^-1^. The simulation was run until steady-state flow conditions were reached. Data analysis excluded this period of increase to only include steady-state flow conditions.

Flow was simulated on a scaled-up version of a real sensilla array and dynamically scaled by matching the Reynolds numbers (*Re*, eq. 1) of the sensilla array based on the sensilla diameter l and flow speed *U* = *U_x_*. To alter the *Re* of each simulation, the fluid density ρ was held constant at 1,000 kg m^-3^ and the dynamic viscosity *μ* of the fluid was changed, varying between 0.122 – 5.50 Pa s to create a range of *Re* between 0.11 – 4.9. Table 1 outlines the parameters used in the simulations. Additional details regarding sampling the parameter space of *Re, Gw, θ* are below in section 2.2.

**Table 1:** Parameters used in simulating fluid velocity fields with constraint-method immersed body method (cIB).

| Parameter | Value range |
| --- | --- |
| Flow speed, $U_x$ (m s<sup>-1</sup>) | 0.06 |
| Reynolds number, $Re$ | 0.11 – 4.9 |
| Distance of array from antenna (m) | 0.02 |
| Diameter of sensillum, $l$ (m) | 0.01 |
| Diameter of antenna (m) | 0.1 |
| Dynamic viscosity of fluid, $\mu$ (Pa s) | 0.122 – 5.50 |
| Density of fluid, $\rho$ (kg m<sup>-3</sup>) | 1000 |
| Angle of array<br>to flow direction, $\theta$ (degrees) | 3.57 – 176 |
| Gap-to-sensillum-diameter ratio, $Gw$ | 1.4 – 49 |
| Space between grid points (m) | $4.88 \times 10^{-4}$ |
| Time step (s) | $1.0 \times 10^{-6}$ |
| Duration of simulation (s) | 0.025 |

#### 2.1.2 Odor concentration model

The velocity of the flow from the immersed boundary simulations were then coupled with an advection-diffusion solver for an odor concentration to measure how much concentration would be absorbed by each array. This model was developed and presented in Waldrop et al (2016).

The concentration, *C*(*x,y,t*) is solved for using

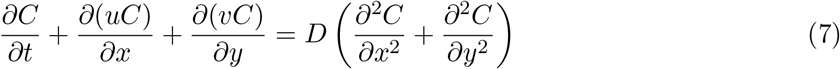

where u = (*u,v*) is the velocity field from the simulations in Section 2.1.1 and *D* is the diffusion coefficient. Equation 7 is solved numerically in a rectangular domain of 1.24m × 1.24m in air and in a domain of 1.25m × 1.25m in water. This is smaller than the domain used to solve for the fluid flow allowing the simulations to be less computationally expensive.

Two different initial odor profiles were used to initialize every simulation. When simulating arrays in water an initial concentration of

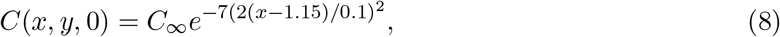

is set where *x* ranges from 1.1 m to 1.2 m (in Waldrop et al (2016), this initial condition is referred to as a thin filament, here with a width of 0.1m). The total amount of chemical present is controlled (integration in x and y on the domain) by setting the maximum value of the concentration, 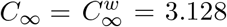. When simulating arrays in air an initial condition, denoted the thick filament, is a never-ending filament. This initial odor concentration has the same exponential profile, Eq. 8 as in the first condition from 1.1m to 1.15m but with 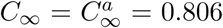 and then 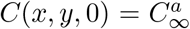 for *x* > 1.15m. The maximum concentrations were set such that the total concentration introduced into each domain is equal during the time of simulation. We allow each simulation to run to time 20 s.

As mentioned above, flow around the array was simulated on a scaled-up version of the array. Therefore, it was necessary to also scale the *Pe* to match the relative rate of mass transport due to advection and diffusion. To do this, we multiplied the original diffusion coefficients from Waldrop et al (2016) by a factor of 2,000. The duration of the simulation was set to 20 seconds to allow the odor filament to penetrate the entire domain. The values used for these simulations are listed in Table 2.

**Table 2:** Parameters used in simulating advection and diffusion of odorant concentration to the sensillum array using velocity vector fields from cIB.

| Parameter | Water simulations | Air simulations |
| --- | --- | --- |
| Diffusion coefficient, $D$ ($\text{m}^2\text{s}^{-1}$) | $1.568 \times 10^{-6}$ | $1.20 \times 10^{-2}$ |
| Grid resolution | 2048 | 512 |
| Width of filament (m) | 0.1 | ∞ |
| Grid size, $h$ (m) | $6.1035 \times 10^{-4}$ | $2.4 \times 10^{-3}$ |
| Time step, $dt$ (s) | $3.7 \times 10^{-3}$ | $5.191 \times 10^{-4}$ |
| Simulation duration, (s) | 20 | 20 |

#### 2.1.3 Numerical Methods

The numerical method used to solve this mathematical model is given in detail in the supplementary information of Waldrop et al (2016) and was implemented in MATLAB. Strang splitting was used to solve the partial differential equation, Equation 7, in multiple steps. Each step was then solved using finite different methods. Here we summarize this method briefly. The following steps are used to advance one timestep:

1. Advection of the concentration for a half timestep:

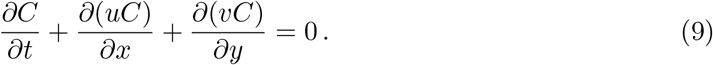 A third-order weighted essentially nonoscillatory method (WENO) is used to solve this step (Shu, 1997).
2. Diffusion of the concentration for a full timestep,

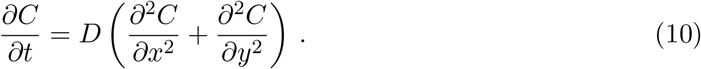 The second order two-dimensional Crank-Nicolson method (Strikwerda, 2004; LeVeque, 2007) is used to solve this step.
3. Determine how much concentration reached each grid point within a sensillum and the concentration was set to 0 at that grid point.
4. Advection of the concentration for another half timestep:

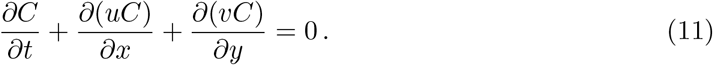

A combination of Dirichlet and no-flux boundary conditions are used in the steps given above. These are set exactly as described in Waldrop et al (2016).

An extensive convergence study was presented in Waldrop et al (2016). To verify that the slight modifications made to method here did not change the convergence behavior of the method, convergence was verified again using a velocity field from Section 2.1.1. Based on these tests we used spatial grids of 512 for the air cases and 2048 for the water cases resulting in a gridsizes reported in Table 2 for the two different conditions. The velocity fields from the immersed boundary simulations in Section 2.1.1 were interpolated to these grids to be able to be used in solving of the concentration. The timestep, *dt*, was set as the smaller of the constraint set by the Courant-Friedrichs-Lewy condition (0.9h/*U*_∞_ where *U*_∞_ = 0.15m s^-1^ is the maximum velocity in all simulations from Section 2.1.1 and h is the spatial grid size) or the constraint set by the diffusive length scale (*R*^2^/4D where *R* = L/2 = 0.005m is the radius of the sensillum and D is the diffusion coefficient) (LeVeque, 2007).

#### 2.1.4 Data analysis

Fluid velocity fields simulated in cIB (example in Fig. 2B) were used to calculate several values related to fluid flow through and around the sensillum array using VisIt (Childs et al, 2012) and *R* statistical software (Team, 2011). Velocity fields used for these calculations are the final time step of the cIB simulation after steady-state flow had been reached. The spatially averaged value of the magnitude of velocities was calculated for a circle of radius 0.01 m around each sensillum: the center sensillum in the array, the ‘top’ sensillum (which represents the sensillum counter-clockwise from the center sensillum), and the ‘bottom’ sensillum (which represents the sensillum clockwise from the center sensillum). The magnitude of velocity was then non-dimensionalized by multiplying by the velocity and the duration of the advection-diffusion simulation time (20 s) and divided by the sensillum diameter (*l* = 0.01 m). All velocities reported are dimensionless.

The shear rate of fluid at the surface of each sensillum was calculated by sampling velocities along a line between the sensilla (see Fig. 2 line labeled *Gw*). Shear rates were calculated at the surface of the sensillum to 30% of the sensillum’s diameter away from the sensillum’s surface (0.003 m). These shear rates are reported for the inside edges of the outer sensilla in the array and the upper edge of the center sensillum in the array. Shear rates were non-dimensionalized by multiplying each shear rate by the advection-diffusion simulation time (20 s). All shear rates reported are dimensionless.

Fluid velocity fields simulated with cIB were also used to calculate the leakiness of the array, defined as the area that fluid that moved through the array in simulation time divided by the area of fluid that could have moved through the same area if the array were absent. Velocities were evenly sampled along a line through the sensilla array (see Fig. 2 orange line labeled *Gw*) using VisIt. These velocities were multiplied by the duration of the simulation (*t* = 0.025) and the distance between points. These values were summed to give the area the fluid travelled through in the simulation. Similarly, the area was then computed using velocity equal to the fixed speed of the simulated flow (0.06 m s^-1^).

Concentration captured by each sensillum during each time step in the advection-diffusion model were summed across the sensilla and temporally to find a total concentration captured value for each simulation. This value was divided by the maximum concentration, *C*_∞_, in each simulation to find the standardized concentration value presented as odor-capture performance.

#### 2.1.5 Computational Environment

Computational simulations were performed on the Bridges Regular Memory cluster at Pittsburgh Supercomputing Center through the Extreme Science and Engineering Discovery Environment (XSEDE) and the Multi-Environment Research Computer for Exploration and Discovery (MERCED) high-performance computing cluster at UC Merced. Bridges Regular Memory is a cluster of 752 nodes with 128GB sRAM each running 2.3 – 3.3 GHz Intel Haswell CPUs with 14 cores each. Data analyses were performed on a Mac Desktop running on 2.7 GHz 12-Core Intel Xeon E5 processor with 64 GB of RAM.

### 2.2 Uncertainty Analysis

In this work, we consider the uncertainty in the following three input parameters, which are represented using uniform distributions: Angle (of sensilla array with respect to oncoming flow, *θ*) 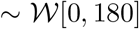 degrees; Gap-width-to-sensillum-diameter ratio (between sensilla to the sensillum diameter, *Gw*) 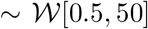; and *Re* (Reynolds number of sensilla array, eq. 1) ~ W[0.01, 5]. To efficiently quantify the uncertainty in the quantities of our interest and analyze the sensitivity of the output quantities with respect to each of the uncertain inputs, we introduce the generalized polynomial chaos (gPC) expansion method to approximate the full simulation and a variance-based sensitivity analysis measure – Sobol indices (SI) – to identify the “importance” of each input.

#### 2.2.1 Generalized Polynomial Chaos Expansion

For each quantity of our interest (denoted as *w*), we construct an approximation *w_p_* with respect to the vector of three uncertain inputs (denoted as *ξ*) using gPC expansion up to order *p* as follows (Wiener, 1938).

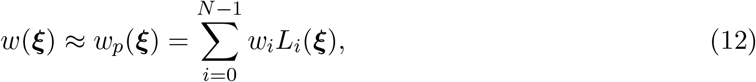

where 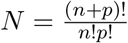 is the number of terms with *n* = 3 as the dimension of inputs, and the parameters *w_i_*s are called the gPC coefficients to be determined. Based on the specific type of distribution the input variables have, one can choose a most proper polynomial basis function from Askey scheme to reach a fast convergence (Xiu and Karniadakis, 2002). In the current work, the functions *L_i_*s are chosen as Legendre polynomials since we consider inputs *ξ* as uniform random variables. The first few univariate Legendre polynomials are

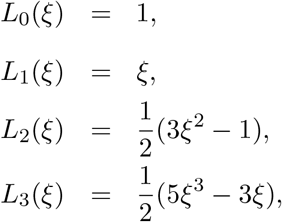

The multivariate Legendre polynomials are the product of univariate polynomials.

To determine the gPC coefficients, we run *M* = 1233 full simulations and extract a set of quantities of interest corresponding to the inputs as 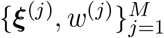, then solve the Least Squares problems for the coefficient vector w = [*w*_1_, *w*_2_,…, *w_N_*] as

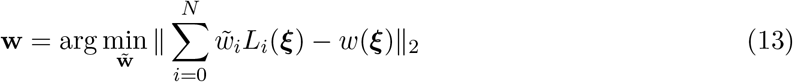

where 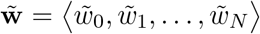 is an arbitrary gPC coefficient vector which converges to the desired coefficient vector w through the minimization.

#### 2.2.2 Sensitivity Analysis

Global sensitivity analysis explores the impact on the model output based on the uncertainty of the input variables over the whole stochastic input space, and it can help to identify the “important” uncertain variables. Here, we adopt a variance-based measure to analyze the global sensitivity analysis: the Sobol indices, which are calculated based on the ANOVA (analysis of variance) decomposition as follows (Sobol, 1993, 2001).

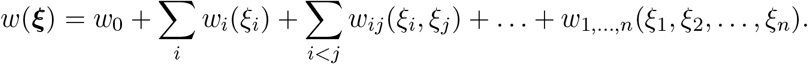

where

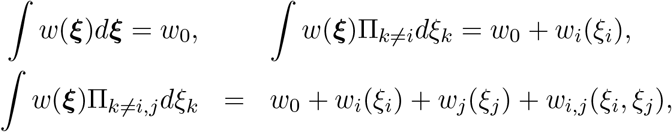

and so on.

Based on the ANOVA decomposition, the variance of the sub-function *w*_*i*_1_,*i*_2_,…,*i*_*r*__ can be defined as

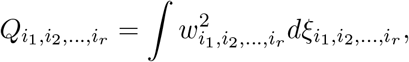

and the total variance is defined as

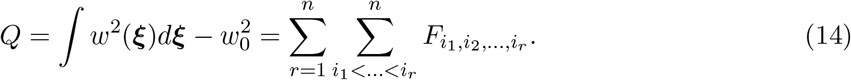

Following which, the global sensitivity indices are defined as the ratio of the variance in sub-dimensional problem to the total variance of the full-dimensional problem as

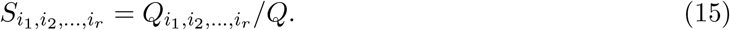

The larger the Sobol index is, the more important the set of input parameters in that sub-dimensional space is. The most frequently used indices are the first order indices and the total indices

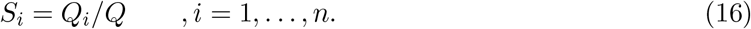

The first order indices measure the sensitivity of the quantity of interest to each single variable *ξ_i_* alone. It can help to rank the “importance” of the variables.

Numerically, one may calculate the Sobol indices using Monte Carlo (MC) method. However, it could be computationally expensive since a large number of full computational fluid dynamics simulations need to be implemented to reach a reasonable convergence. Therefore, we calculate the Sobol indices based on the gPC expansion in this work (Sudret, 2008). Assume the gPC expansion is obtained as

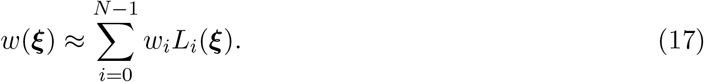

The multivariate Legendre polynomial *L_i_* can be represent by products of univariate polynomial with multiple index *α* = (*α*_1_,…, *α_n_*) as

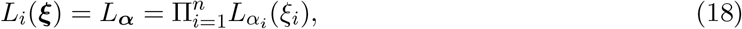

Let 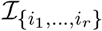 denote the set of *α* multi-indices where only *α_k_* ≠ 0 for *k* = *i*_1_, *i*_2_,…, *i_r_*. Then the gPC expansion can be rewritten as

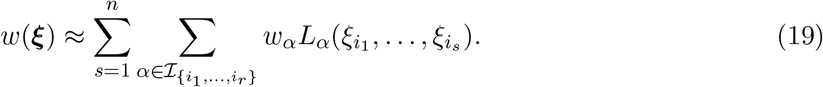

Based on that, the Sobol indices can be approximated using,

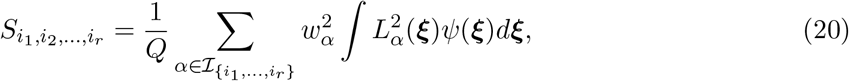

where *ψ(ʾ)* is the probability density function of 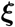 and

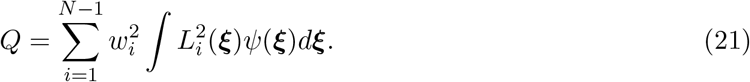

The set of Sobol indices is a variance-based measure to analyze the sensitivity of the model output (quantity of interest) to each single variable and the sets of variables. SIs of all the quantities of interest sum to 1 and they shows that the variation of a specific variable or a specific set of variables makes the majority contribution to the output variance. By comparing the Sobol indices, one can rank the importance of the uncertain variables and focus on the exploration of those important variables in the physical process.

### 2.3 Public Data Availability

Raw data and the code to produce all data figures are publicly available at figshare: 10.6084/m9.figshare.6399740, 10.6084/m9.figshare.6399743, 10.6084/m9.figshare.6399746, and 10.6084/m9.figshare.6399749.

## 3 Results

### 3.1 Flow in the array

Fig. 2B rep re se nts a typical velocity vector field from the last time step of the advection simulation in cIB (*Re* = 0.24, *Gw* = 6.08, *θ* = 38). Flow processes around the antenna and the sensilla array, flow being much slower for sensilla that; are either directly upstream or downstream of the antenna.

Fig. 3 reports values for average speed (A – C) around each sensillum and Fig. 4A and Table 3 report values of Sobol indices. The orientation of each sensillum to the antenna changes with the angle *θ*, and average speed around each sensillum is sensitive to *θ* (center sensillum SI = 0.966, top sensillum SI = 0.260, bottom sensillum SI = 0.252). Average speed around the center sensillum, the closest sensillum to the antenna, is especially sensitive to changes in *θ* since the antenna often shadows the center sensillum at very low and very high values of *θ* (Fig. 3A). The average fluid speeds around the two outer sensilla seem most sensitive to changes in *Re* (top sensillum SI = 0.400, bottom sensillum SI = 0.420) and not *Gw* (top sensillum SI = 4.35 ×10^−4^, bottom sensillum SI = 7.51 ×10^−3^). Averaged speed around these sensilla is significantly influenced by the interaction between *θ* and *Gw* (top sensillum SI = 0.332, bottom sensillum SI = 0.324), supporting previous studies on flow through sensilla arrays.

**Figure 3:**
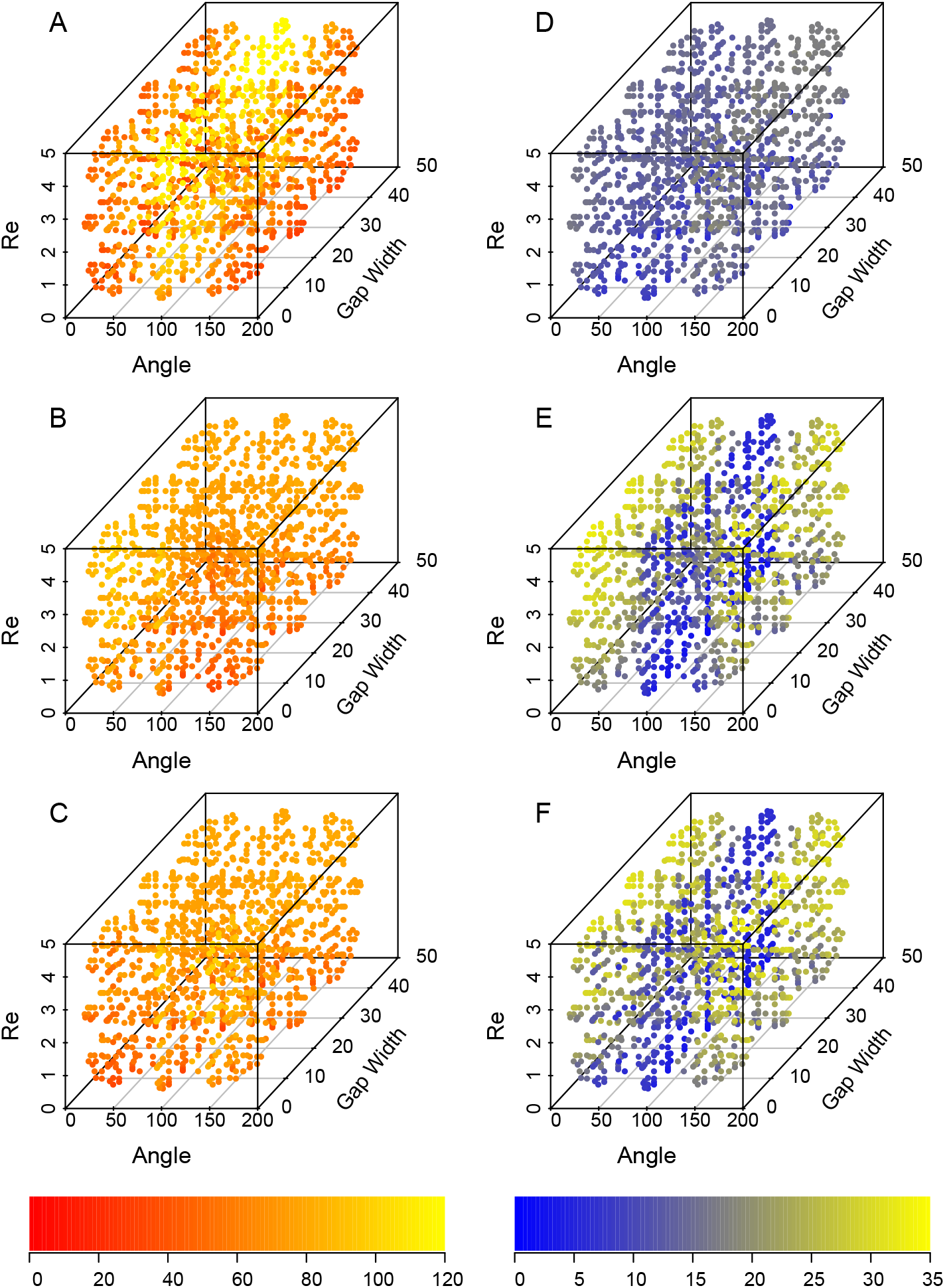
Mean magnitudes of dimensionless velocity (A – C) and dimensionless shear rates (D – F) against each parameter (*θ, Gw, Re*) for each olfactory sensillum in the array. A,D: center sensillum, B,E: top sensillum (counter-clockwise from center sensillum), C,F: bottom sensillum (clockwise from center sensillum). Color scale at bottom indicates mean magnitude of velocity (left) shear rate (right) from lowest to highest on each plot.

**Table 3:** Sobol indices (SI) calculated for each of the three parameters (*Re, Gw, θ*) and their interactions for average speed, shear rate, leakiness, concentration captured in water, and concentration captured in air. SI’s for each sensillum are reported on each line of the columns for average speed and shear rate (first line: center sensillum; second line: top sensillum; third line: bottom sensillum).

|  | Average speed | Shear rate | Leakiness | Conc. captured<br>in water | Conc. captured<br>in air |
| --- | --- | --- | --- | --- | --- |
| $Re$ | 0.138,<br>0.736,<br>0.736 | 0.489,<br>0.118,<br>0.116 | 0.862 | 0.880 | $2.37 \times 10^{-5}$ |
| $Gw$ | $5.16 \times 10^{-5}$,<br>$4.19 \times 10^{-3}$,<br>$4.22 \times 10^{-3}$ | $4.76 \times 10^{-3}$,<br>$4.87 \times 10^{-3}$,<br>$6.06 \times 10^{-3}$ | 0.0151 | $5.20 \times 10^{-4}$ | 0.833 |
| $\theta$ | 0.8518,<br>0.110,<br>0.110 | 0.498,<br>0.849,<br>0.845 | 0.0686 | 0.114 | 0.160 |
| $Re \& Gw$ | $1.66 \times 10^{-5}$,<br>$7.87 \times 10^{-5}$,<br>$7.83 \times 10^{-5}$ | $2.71 \times 10^{-3}$,<br>$6.69 \times 10^{-5}$,<br>$9.70 \times 10^{-5}$ | $6.241 \times 10^{-4}$ | $6.14 \times 10^{-5}$ | $1.68 \times 10^{-6}$ |
| $Re \& \theta$ | $8.82 \times 10^{-3}$,<br>$7.46 \times 10^{-4}$,<br>$7.51 \times 10^{-4}$ | $3.52 \times 10^{-3}$,<br>0.0143,<br>0.0112 | 0.0214 | $4.34 \times 10^{-3}$ | $3.79 \times 10^{-6}$ |
| $Gw \& \theta$ | $4.41 \times 10^{-4}$,<br>0.149,<br>0.149 | $4.59 \times 10^{-3}$,<br>0.0134,<br>0.0223 | 0.0317 | $2.75 \times 10^{-4}$ | $7.05 \times 10^{-3}$ |
| All<br>parameters | $2.17 \times 10^{-9}$,<br>$7.89 \times 10^{-5}$,<br>$7.88 \times 10^{-4}$ | $7.84 \times 10^{-8}$,<br>$1.97 \times 10^{-5}$,<br>$1.61 \times 10^{-4}$ | $5.20 \times 10^{-4}$ | $3.68 \times 10^{-6}$ | $3.34 \times 10^{-11}$ |

Shear rates (Fig. 3 D – F) were absolutely much higher for the outer sensilla in the array compared to the center sensillum. Shear rates for all sensilla are highly sensitive to *θ* (center sensillum SI = 0.498, top sensillum SI = 0.849, bottom sensillum SI = 0.845; Fig. 4B) and moderately sensitive to *Re* (center sensillum SI = 0.489, top sensillum SI = 0.118, bottom sensillum SI = 0.116). The shear rates around the center sensillum are more heavily influenced by *Re* mostly likely due to its close proximity to the antenna. *Gw* does not influence shear rates on any of the sensilla (center sensillum SI = 4.76 × 10^−3^, top sensillum SI = 4.87 × 10^−3^, bottom sensillum = 6.06 × 10^−3^), nor do any interactions between parameters (all interaction SI’s < 0.03).

**Figure 4:**
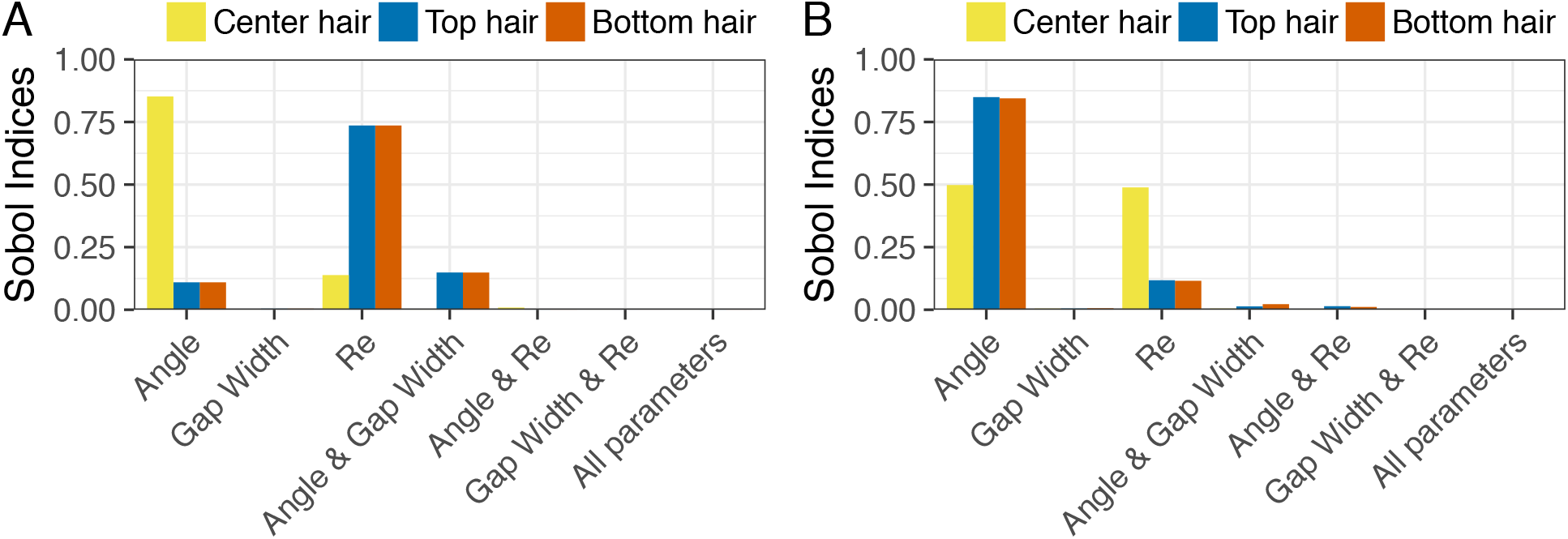
Sobol indices calculated for each parameter *Re*, *Gw*, and *θ* as well as interactions between parameters. A: Average speed and B: shear rate. Each bar represents a sensillum in the array.

Values of leakiness ranged from 2.56×10^−5^ to 0.0317, much lower than previous studies due to the influence of the central antenna which diverted much of the flow around the sensilla array at high and low values of *θ*. The sensitivity of leakiness (Fig. 5C and D, purple bars) of the array was dominated primarily by changes in *Re* (SI = 0.862) and to a much lesser extent *θ* (SI = 0.0686). Notably, *Gw* and interactions between *Re* and *Gw* did not have a major influence on leakiness (SI = 0.0151 and SI = 6.24×10^−4^, respectively), contrary to previous studies of leakiness in sensilla arrays (Cheer and Koehl, 1987a,b). This is likely due to the influence of the central antenna which diverted flow around the array at many of the values of *Gw* that would otherwise produce higher values of leakiness.

**Figure 5:**
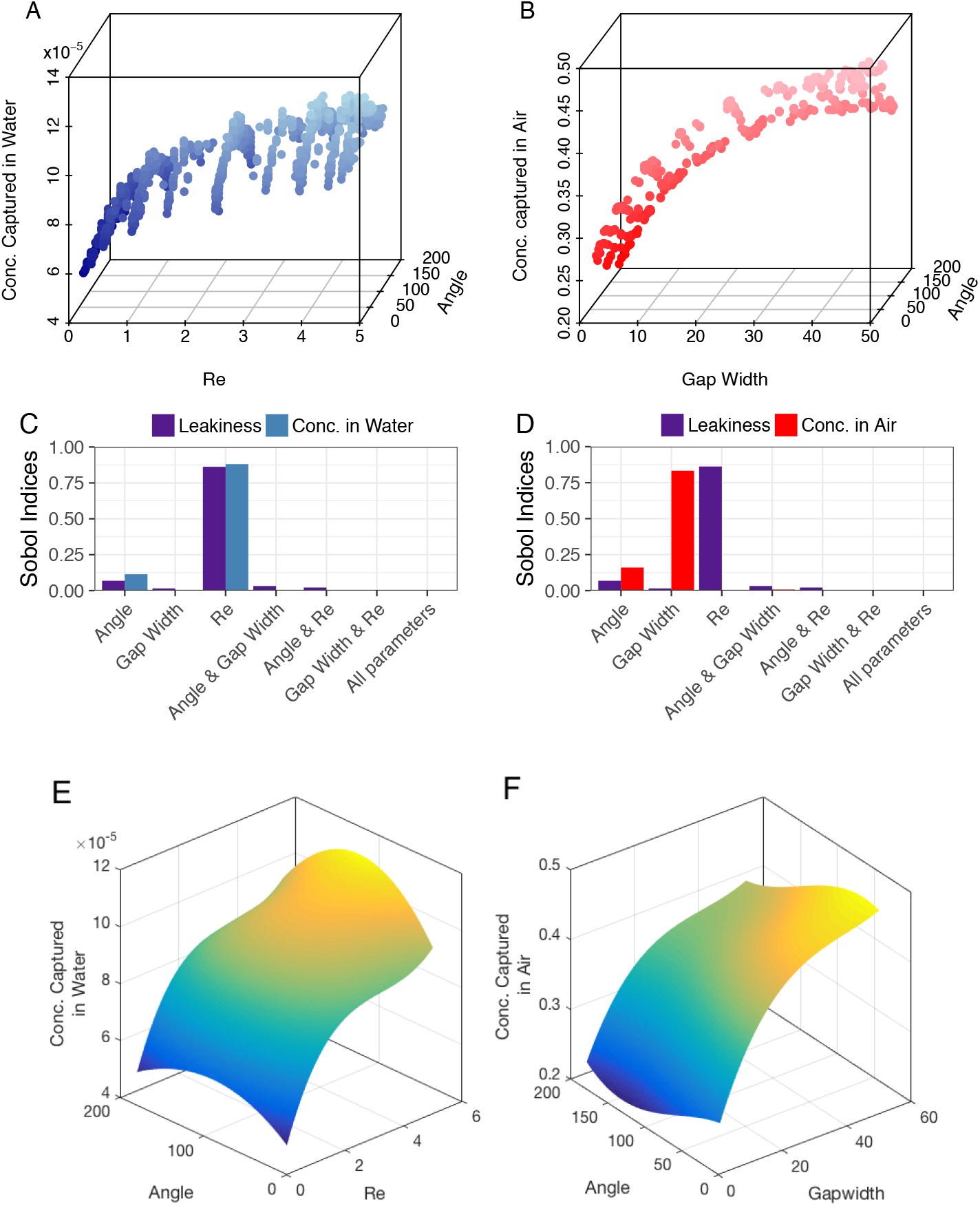
Concentration captured versus select parameters and Sobol indices. A: Concentration captured in water versus angle *θ* and *Re*. C: Sobol indices are reported for leakiness (purple) and concentration captured in water (blue) for comparison. B: Concentration captured in air versus gap-width-to-diameter ratio (*Gw*) and angle *θ*. D: Sobol indices are reported for leakiness (purple) and concentration captured in air (red) for comparison. Surrogate plots calculated for concentration captured in water (E) and in air (F) with two parameters in top row.

### 3.2 Odor capture in water and air

Fig. 5 reports values of standardized concentration of odor captured in water (left column) and in air (right column) conditions. Concentration captured in water ranged from 5.56×10^−5^ to 1.20×10^−4^, while concentration captured in air ranged from 0.223 to 0.489. The difference in performance is reflective of absolute differences of capture between both air and water.

The sensitivities of concentration captured to parameter change differed dramatically between water and air conditions (Fig. 5C and D, respectively). Sobol indices indicate that capturing odor concentration in water is dominated primarily by *Re* (SI = 0.880) and to a lesser extent *θ* (SI = 0.114), being relatively insensitive to changes in *Gw* (SI = 5.20×10^−4^) and interactions between parameters (SI’s < 0.01). The surrogate plot of concentration captured in water versus *Re* and *θ* (Fig. 5E) shows that high *Re* lead to higher capture in water, and middling values of *θ* enhance the high values of *Re* (when the sensilla array is on the upper side of the antennule to flow, not being shadowed by it).

In contrast, concentration captured in air is highly sensitive to *Gw* (SI = 0.833) and moderately sensitive to *θ* (SI = 0.160), while being insensitive to *Re* (SI = 2.37×10^−5^) and interactions between parameters (SI’s < 0.001). This is a reversal of the trend of concentration capture in water which shows low sensitivity to *Gw* and high sensitivity to *Re*. The surrogate plot of concentration captured in air versus *Gw* and *θ* indicates that the highest captures occur at high *Gw* values (when sensilla are spaced far apart) and low *θ* values (when the sensilla array is on the upstream side of the antenna).

Performance in the context of parameter change are similar between leakiness and concentration captured in water and dissimilar between leakiness and concentration captured in air. Sobol indices reported for the three performance metrics in Fig. 5C and D reflect this pattern: leakiness and concentration captured in water share similar values for sensitivity to *Re* (0.862 and 0.85, respectively) and *Gw* (0.0151 and 6.60×10^−4^, respectively), while leakiness and concentration captured in air show drastically different Sobol indices for the two parameters. The norm of the difference between values of leakiness and concentration captured in water is much lower (0.313) than the same norm calculated between values of leakiness and concentration captured in air (0.445), indicating that leakiness is a better predictor of concentration captured in water than concentration captured in air.

## 4 Discussion

### 4.1 Odor capture differs between air and water

In this study, we simulate odor capture by a series of theoretical olfactory sensilla arrays in two biologically and environmentally relevant situations reflective of water and air, respectively: a narrow, brief band of high-odor concentration and a broad, continuous band of lower-odor concentration. We use this model to investigate the effects of three parameters (Reynolds number *Re*, Gap-to-sensillum-diameter width Gw, and the angle of the array to the direction of oncoming flow *θ*) on odor-capture performance. Leakiness and concentration captured values provide a way to assess the relative performance of individual sensillum arrays in a typical encounter event in water and in air. Sobol indices calculated via uncertainty analysis provide a quantitative way to assess the relative sensitivity of capture to the parameters varied.

The results of odor-capture performance indicate that there are profound differences between odor capture in air versus water. Many times more odorant is captured in air (Fig. 5A-B,E-F)), likely the result of the longer exposure times to a larger filament which is many times the diameter of the olfactory sensillum and the higher diffusivity of odorants in air compared to water (Willis and Arbas, 1991). The coefficient of diffusion used for these simulations were 10,000 times higher in air than in water, resulting in low Peclet numbers (*Pe* < 1) and facilitating greater capture in air. This result held even though the odor filament in air was of lower concentration than the high-concentration filament of water, which is broadly reflective of odor filaments in their respective fluid environments.

Furthermore, the sensitivity analyses on odor capture revealed that the parameters of the array controlling performance are drastically different between air and water. The quantitative sensitivity analysis indicates that gap width between sensilla *Gw* has no meaningful effect on odor capture in water (Fig. 5D), a result seemingly contrary to previous studies (Cheer and Koehl, 1987a,b; Stacey et al, 2002; Schuech et al, 2012). The presence of the antenna greatly reduces leakiness of the array compared to previous studies, consistent with other studies investigating leakiness through a sensillum array close to solid boundaries (Loudon et al, 1994). *Re* has an important effect on odor capture in water (Fig. 5D) as low diffusion rates of odorant molecules leads to the boundary layer being more of a barrier for arrays in water rather than air. Boundary layers share inverse relationship with *Re* and both leakiness and odor capture in water are highly sensitive to changes in *Re*.

Capture in air is tied more to sensillum arrangement than the size or movement of the array. In contrast to both odor capture in water and leakiness, *Re* does not influence odor-capture performance in air (Fig. 5D). Instead, the arrangement of sensilla, i.e. the gaps between sensilla *Gw* and to a lesser extent *θ*, seem to influence odor capture in air. This is likely a reflection of the outsized influence of diffusivity to capture at low *Pe*, and where spreading sensory sensilla further apart increases the area swept through and avoids the slow-moving fluid around the antenna.

Leakiness and other measures of fluid flow are more relevant predictor of odor capture performance in water than in air. Many studies have assumed leakiness as a proxy for odor-capture performance in aquatic crustaceans (Mead et al, 1999; Mead and Koehl, 2000; Humphrey and Mellon, 2007; Reidenbach et al, 2008; Nelson et al, 2013; Waldrop et al, 2015b,a), and the results of this study generally support this assumption in water where *Pe* are high (on the order of 1,000). However, the dramatic differences in capture in air do not support this assumption; diffusion rates D in air are high enough to dominate capture, having *Pe* below one. Studies that have investigated both array leakiness and odor-capture performance have found a similar mismatch between the two (Waldrop and Koehl, 2016; Waldrop et al, 2016). Diffusive transport must be taken into account when investigating capture by sensillum arrays in air.

### 4.2 Predictions of Morphological Diversity

Animals are under different sets of constraints capturing odors in water versus air. Fluids possess different physical properties, diffusion rates vary dramatically between fluids, and the shape, size, and frequency of the odor filaments in the environment all differ as a result of fluid habitat. Thus, it is reasonable to expect differences in common combinations of morphological parameters for extant groups of animals in aquatic versus terrestrial habitats.

Most biological systems are extremely complex, and odor capture is no exception. The complexity of parameter space leads to many parameter combinations that will result in the same functional performance such as many-to-one mapping, mechanical equivalence, or functional redundancy (Wainwright et al, 2005; Anderson and Patek, 2015; Muñoz et al, 2017). Since odor capture is a functionally redundant performance metric, making specific predictions about the optimal configuration of parameters involved in odor capture is less useful than making broader predictions about the ways in which the parameters of this system could be constrained, the rates at which diversification could be expected, and the resulting morphological range of parameter values. Here, we make some general predictions on the diversity of parameter values in groups of aquatic and terrestrial animals within the Pancrustacea based on the performance results of our model and sensitivity analyses.

Performance in aquatic crustaceans should be more constrained overall due to lower capture rates and the nature of odor filaments in water. Our simple model shows that performance in water is tied heavily to Reynolds number (Re), but not factors associated with the arrangement of the sensilla. Since odor-capture performance in water is highly sensitive to *Re* but less so to Gw, it is likely that aquatic crustaceans will have more dense arrays that rely on animal-generated currents to increase *Re* fluid penetration into the array. Additionally, the range of *Re* over which aquatic crustaceans operate should be constrained and show low diversity or be the sites of potential rapid evolutionary change between antennule configurations which allow for high odor-capture performance.

Conversely, performance in terrestrial crustaceans should not be directly tied to changes in flow (*U* in *Re* eq. 1) caused by either animal-induced or environmental fluid currents, but should show higher constraints in terms of the gaps between olfactory sensilla. The simple model shows that performance in air is likely unconstrained by Re, and instead may rely heavily on gap-width-to-sensillum-diameter ratio (*Gw*), spacing olfactory sensilla far apart. Terrestrial crustaceans, including insects, should have arrays that are relatively sparse compared to aquatic crustaceans and wider morphological diversity. Anecdotally, this seems to be true given studies on insects which exhibit a wide diversity of array and olfactory sensilla morphologies (Hallberg and Hansson, 1999).

In the case of animals that require specialized sensilla that pick up very rare or dilute signals (such as sex pheromones) may require insects to maintain denser array with smaller gap widths between sensilla in increase the overall surface area of the sensor. When arrays are very dense and sensilla close together, it is possible that animals would be required to induce flow through flapping, waiving, or flying, where low leakiness and the local *Pe* would be dominated by advective transport and odor capture would be more sensitive to manipulation of *Re* of the array. This idea is supported anecdotally by observations of terrestrial hermit crabs that have dense arrays with very small gaps that lay close to the antennule and flick their antennules (Waldrop and Koehl, 2016; Waldrop et al, 2016) and silkworm moths that have extremely dense arrays and require wing-flapping to generate currents within the array to aid in odorant capture (Loudon et al, 2000). In these cases, increased flow within the dense arrays may make available more of the sensory surface area than no animal-generated flow (Waldrop et al, 2016).

### 4.3 Future directions

Uncertainty analysis on computational models provides a quantitative way to analyze the relative importance of morphological and kinematic parameters on functional performance. It provides an important bridge between computational modeling and studies of morphological diversity in animal populations, making it possible to predict patterns of morphological and kinematic diversity based on the sensitivity of each parameter on performance.

However, these predictions are only as good as the model itself at reflecting the most important parameters of each functional system. Our model accounts for only three of many parameters that may be influential to odor capture in two dimensions due to computational constraints. The model excludes many parameters that could have a major effect on performance and thereby change the predictions made by sensitivity analyses.

As a result, these hypotheses need to be tested against morphological and kinematic measurements of several groups of aquatic and terrestrial crustaceans representing independent lineages of terrestrialization. Phylogenetically corrected measurements of performance can be used to test predictions about the expected rates of parameter evolution and the overall diversity of parameter values in groups (Muñoz et al, 2017). Additionally, measurements of these parameters on groups of aquatic and terrestrial crustaceans can be measured to see if parameters are constrained or diverse compared to predictions made by the sensitivity analyses.

While a very simple model with many limitations, morphological measurements from extant animals can provide important feedback for improving the model and better understanding odor capture. Comparison to a systematic range of extant odor-capture antennae will provide important feedback to the model and point towards parameters that should be included to refine the predictions made by sensitivity analyses through other tools in uncertainty quantification.

## Acknowledgements

The authors wish to acknowledge the following funding sources: funds from the New Mexico Institute of Mining and Technology to L. Waldrop; computational allocations to L. Waldrop from the Extreme Scientific and Engineering Discovery Environment (XSEDE) TG-CDA160015 and TG-BI0 170090; and funding to S. Khatri from the National Science Foundation Physics of Living Systems #1505061.

The authors wish to thank Swayamjit Ray, Anjel Helms, and Loren Rivera for organizing the “Chemical Ecology in the New Era of Technology” symposium; Laura Miller, Amneet B., and Boyce Griffith for help with IBAMR; David O’Neal at Pittsburgh Supercomputing Center for aid in computing; and Sheila Patek, Philip Anderson, Jonathan Rader, and Dennis Evangelista for influential discussions regarding evolutionary biomechanics.

